# SBpipe: a collection of pipelines for automating repetitive simulation and analysis tasks

**DOI:** 10.1101/107250

**Authors:** Piero Dalle Pezze, Nicolas Le Novère

## Abstract

**Background:** The rapid growth of the number of mathematical models in Systems Biology fostered the development of many tools to simulate and analyse them. The reliability and precision of these tasks often depend on multiple repetitions and they can be optimised if executed as pipelines. In addition, new formal analyses can be performed on these repeat sequences, revealing important insights about the accuracy of model predictions.

**Results:** Here we introduce SBpipe, an open source software tool for automating repetitive tasks in model building and simulation. Using basic configuration files, SBpipe builds a sequence of repeated model simulations or parameter estimations, performs analyses from this generated sequence, and finally generates a LaTeX/PDF report. The parameter estimation pipeline offers analyses of parameter profile likelihood and parameter correlation using samples from the computed estimates. Specific pipelines for scanning of one or two model parameters at the same time are also provided. Pipelines can run on multicore computers, Sun Grid Engine (SGE), or Load Sharing Facility (LSF) clusters, speeding up the processes of model building and simulation. SBpipe can execute models implemented in Copasi, Python or coded in any other programming language using Python as a wrapper module. Future support for other software simulators can be dynamically added without affecting the current implementation.

**Conclusions:** SBpipe allows users to automatically repeat the tasks of model simulation and parameter estimation, and extract robustness information from these repeat sequences in a solid and consistent manner, facilitating model development and analysis. The source code and documentation of this project are freely available at the web site: https://pdp10.github.io/sbpipe/.

## Background

The range of software tools developed by the Systems Biology community has grown considerably in the last few years, in particular aimed at supporting mathematical modelling of biological networks. The development of a mathematical model typically comprises successive phases: design, parameterisation, simulation and testing. Model design is the phase where the core of the problem to investigate is summarised using a mathematical formalism. Once designed, the model parameters need to be calibrated, for example using some experimental data. After this stage, the model is used for generating predictions which are then tested experimentally. Depending on the outcome, a model can be refined in order to improve or correct its prediction. Many tools already exist to generate, simulate and analyse mathematical models [1,2]. Although these tools provide modellers with key functionalities for model parameter estimation and simulation, it has become clear that the accuracy of these tasks depends on multiple repetitions. Furthermore, the analysis of this batch of repeats can reveal important insights regarding the model itself and the data used for calibration. Therefore, it is useful to repeat tasks such as parameter estimation or stochastic simulation, collect statistics and visualise these results.

SBpipe is an open source software tool which provides modellers with a collection of pipelines for model development and simulation. A pipeline for parameter estimation allows users to repeat a model calibration many times on a multicore machine or a computer cluster. The generated fit sequence is then analysed, and information about the profile likelihood from parameter estimation samples is represented graphically and textually. Support for model simulation is also provided with pipelines for time course model simulation, as well as single and double parameter scans.

## Implementation

SBpipe is an open source software package developed with the Python [3] and R [4] programming languages. Python is the main programming language connecting all the package components, whereas R is used for generating statistics and plots. The use of this statistics-dedicated programming language for analysing the results allows users to run the provided R scripts independently of SBpipe using an R environment. This can be convenient if further data analysis are needed or plots need to be annotated or edited.

Pipelines in SBpipe are configured using configuration files. This allows modellers to easily edit their tasks manually or programmatically if needed. Examples of configuration files can be found within the main package in the folder `tests/insulin receptor/`

In order to maintain a flexible and extendible design, SBpipe abstracts the concepts of simulator and pipeline. The class `Simul` is a generic simulator interface used by the pipelines in SBpipe. This mechanism uncouples pipelines from simulators which can therefore be configured in each pipeline configuration file. Currently, the available simulators are `Copasi` and `Python`. These simulators process models developed in COPASI and models coded in Python, respectively.

SBpipe passes the report file name as an input argument to the latter. The Python program is then responsible for generating a report file containing the simulation (or parameter estimation) results. Python can also be used as a wrapper module for running models coded in any programming language. Indeed instead of coding a model itself, the Python file can call an external program containing the model. This Python wrapper must forward the report file name to this external program which becomes responsible of generating the report file. With this simple approach, users can run their existing models using customised command options or any program library they need. The `tests/` folder contains examples of models coded in R, Octave, or Java programming languages, and executed using basic Python module wrappers. The supplied R models depend on the packages minpack.lm, deSolve, and sde, whereas the supplied Java model requires a JVM. Dependencies for these additinal models must be installed separately.

The class `Pipeline` represents a generic pipeline, which is extended by each SBpipe pipeline. The following pipelines are currently available:

– `simulate:` deterministic or stochastic time course stimulation;
– `single param scan:` scan a model parameter;
– `double param scan:` scan two model parameters;
– `param_estim:` model parameter estimation including sampling of the parameter likelihood.

An SBpipe pipeline performs three tasks: data generation, data analysis, and report generation. Depending on the configuration file options, the first task loads and runs a simulator at runtime and organises the generated data. The second task computes statistics and plots from these data. Finally, the third task generates a LaTeX/PDF report containing the computed plots.

Pipelines for parameter estimation or stochastic model simulation can be computationally intensive. SBpipe allows users to generate repeats of model simulation or parameter estimation in parallel. In a configuration file, users can select the number of repeats, and whether the jobs should be executed locally (multithreading is available) or in a computer cluster. In this case, SBpipe supports the following cluster types: Sun Grid Engine (SGE) and Load Sharing Facility (LSF).

The project is available on the GitHub repository. Numerous test cases are also provided within the package. Every time the source code is updated online, these tests are automatically executed by Travis.CI, a GitHub application for continous integration service. For standard users, these tests are useful examples of how to configure SBpipe. User and developer documentations for this project are available online and within the project folder.

## Results

A minimal model of the insulin receptor (IR) is used as an example (Additional file 1: Table S1, Figure S1). The generic pipeline work flow is shown for the parameter estimation pipeline in Figure 1A. To illustrate how SBpipe can reveal parameter identifiability issues from multiple parameter estimations, two fit sequences are independently generated using sufficient and insufficient data sets (Additional file 1: Tables S2-S4). For each group, SBpipe generates *N* = 1000 independent parameter estimations using Particle Swarm optimisation algorithm as implemented in Copasi. These calibrations are then processed in the data analysis task. Although SBpipe does not contain a pipeline for computing identifiability analysis directly, the parameter estimation pipeline can help identify issues in parameter estimation by projecting the estimates for each parameter. This analysis uses not only the best fit of each of the *N*estimations, but also the sub-optimal fits. As these fits represent samples of the parameter space, they can reveal a *sampled profile likelihood estimation (PLE)* for each estimated parameter. For direct methods calculating model parameter profile likelihoods using COPASI, see [5] or https://pypi.python.org/pypi/PyCoTools. Results of estimation tasks using data sets presented in Table S2A and Table S2B are shown in the *Identifiable* or *Non-identifiable* columns of Figure 1, respectively.

**Figure 1.**
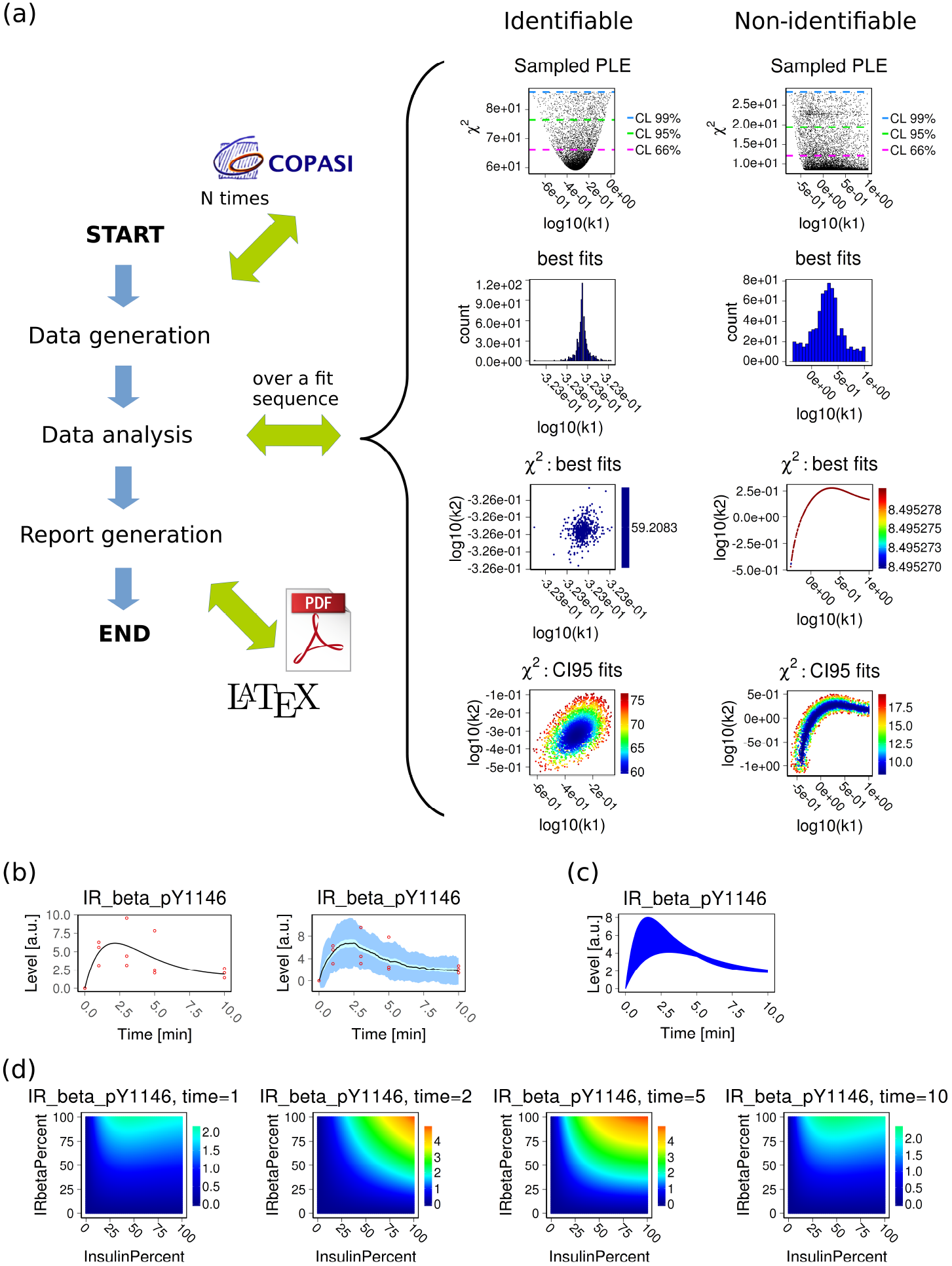
Implemented pipelines in SBpipe. (a) Example of work flow using the parameter estimation pipeline. Parameter estimations were performed using data sets of different sizes. The *Identifiable* column shows the results using a data set sufficient for estimating the parameters with their confidence intervals, whereas the column *Non-identifiable* illustrates the results using the same model but a reduced data set, insufficient for identifying parameter values. Size of the fit sequence: N=1000. For the complete results generated by this pipeline, see Additional file 1: Tables S2-S4, Figures S2-S8. (b) Deterministic and stochastic model time courses for the phosphorylated IR beta species obtained with the model simulation pipeline. For stochastic simulations, mean (black), 95% confidence interval for the mean (cyan), and 1 standard deviation (light blue) are reported. Experimental data are added and indicated as red circles. For the complete results, see Additional file 1: Figures S9-S10. (c) Single parameter scan pipeline. The k1 parameter regulating the IR beta phosphorylation was scanned within its 95% estimated confidence interval. The blue area is the results of 100 time course simulations over this interval. For the complete results, see Additional file 1: Figures S11-S12. (d) Double parameter scan pipeline. Signal intensities for the phosphorylated IR beta receptor different levels of Insulin (x axis) and IR beta receptor (y axis) at 1, 2, 5, and 10 minutes upon insulin stimulation. The colour representation indicates how the readout signal intensity varies upon two model parameter levels. For the complete results, see Additional file 1: Figures S13-S15.

The *Identifiable* column shows how the parameter *k*1 presents clear confidence intervals at 66%, 95%, and 99% percents of confidence levels (CL). The *Non-identifiable* column shows how the same parameter is practically non-identifiable to the right of the confidence interval. Parameter distributions and correlations are also computed for the best fits, and for the fits within each confidence levels. For the complete results generated by this pipeline, see Additional file 1: Tables S2-S4, Figures S2-S8. Results generated by the time-course simulation pipeline are shown in Figure 1B. Deterministic and stochastic model simulations are illustrated for the phosphorylated state of the IR species. For deterministic simulation, time courses of model variables are simply plotted. For stochastic simulations, SBpipe can represent time courses with mean (black line), the 95% confidence intervals of the mean (cyan bars), and one standard deviation (blue bars). The second panel in Figure 1B show this plot using a sequence of 40 independent stochastic simulations. If available, data corresponding to model variables can easily be added to the plot by specifying the data set file name in the configuration file. For the complete results, see Additional file 1: Figures S9-S10.

Figure 1C shows the results from the single parameter scan pipeline. Simulations are ran with values of the parameter `k1` within the 95% confidence interval as determined by the parameter estimation using the data with a sufficient number of data points. If needed, differential scales can also be configured in order to discriminate protein levels. This is particularly useful if a simulated protein knockdown (or overexpression) is investigated. For the complete results, see Additional file 1: Figures S11-S12.

Results generated by the double parameter scan pipeline are shown in Figure 1D. In this analysis two model parameters are scanned simultaneously and these data are reported for each time point separately. For instance, it can be useful for revealing combinatorial effects of two drugs affecting a timecourse. For the complete results, see Additional file 1: Figures S13-S15. An example of this analysis can be found in [6], where it was applied for exploring the combination of mTOR and ROS treatments in a cellular senescence model.

## Discussion

SBpipe is a software tool which allows modellers to automatically repeat certain tasks in model development and analysis, such as parameter estimation and simulation, and obtain additional information about the robustness of the model. Its use should increase productivity and the confidence in the results obtained with the model.

Parameter estimation from experimental data is a challenging task which can easily produce unreliable results due to local minima, parameter non-identifiability, or inadequate optimisation algorithm configuration. From the generation and analysis of a fit sequence, SBpipe can reveal crucial insights about a model structure, the reliability of each parameter, as well as indications about the sufficiency and quality of the experimental data used to calibrate the model. This knowledge is required for assessing whether parameters are well defined and the overall model predictions are reliable.

SBpipe presents advanced pipelines in a simplified manner. Users only need to create a configuration file and run it using a simple command set. A price to pay for this simplicity is that contrary to standard pipeline frameworks, SBpipe does not currently offer support for dependency management at coding level and reentrancy at execution level. The former is defined as a way to precisely define the dependency order of functions. The latter is the capacity of a program to continue from the last interrupted task. Although many pipeline framworks are available for bioinformatics, the definition of a clear and spread standard specifying how pipelines can be configured is still limited in our opinion. In the future we hope to also use a pipeline framework as an additional way to run SBpipe tasks. Benefitting of dependency declaration and execution reentrancy would in particular be beneficial for running SBpipe on clusters or on the cloud.

From an implementation standpoint, SBpipe design is sufficiently generic to permit rapid extension of new pipelines. With this solid but flexible design, SBpipe aims to encourage the development of pipelines for systems modelling into a single community activity.

## Conclusions

SBpipe is a novel open source software that enables systems biology modellers to simulate models, scan and estimate model parameters in a large scale. Novel analyses from multiple repeats are also computed via publication quality plots and tables. This project permits to increase productivity and reliability in model building and simulation.

## Availability and requirements

**Project name:** SBpipe

**Project home page:** https://pdp10.github.io/sbpipe/

**Operating system(s):** Platform independent

**Programming language:** Python 2.7+ or 3.4+, R 3.3.0+

**Other requirements:** Copasi 4.19, TexLive 2013.

**License:** GNU LGPL v3

## List of Abbreviations

CL: Confidence Level IR Insulin Receptor
LSF: Load Sharing Facility
PLE: Profile Likehood Estimation
SGE: Sun Grid Engine

## Author’s contributions

PDP and NLN conceived and designed the project; PDP implemented the software. PDP and NLN wrote the manuscript. All authors read and approved the final manuscript.

## Acknowledgements and Fundings

We acknowledge Dr Lu Li, Dr An Nguyen, and Dr Pınar Pir for helpful feedback. The work was funded by British BBSRC (BBS/E/B/000C0419).

## Additional Files

Additional file 1 — Supporting information

Additional file containing supporting Tables S1-S4 and Figures S1-S15.

## References

1. Ghosh S, Matsuoka Y, Asai Y, Hsin KY, Kitano H. Software for systems biology: from tools to integrated platforms. Nature Reviews Genetics. 2011;12:821–832.

2. Le Novère N. Quantitative and logic modelling of molecular and gene networks. Nature Reviews Genetics. 2015;16:146–158.

3. van Rossum G. Python Programming Language. In: Chase J, Seshan S, editors. USENIX Annual Technical Conference. USENIX; 2007.

4. R Development Core Team. R: A Language and Environment for Statistical Computing. Vienna, Austria; 2008. ISBN 3-900051-07-0.

5. Schaber J. Easy parameter identifiability analysis with COPASI. Biosystems. 2012;110(3):183–5.

6. Dalle Pezze P, Nelson G, Otten E, Korolchuk V, Kirkwood T, von Zglinicki T, et al. Dynamic Modelling of Pathways to Cellular Senescence Reveals Strategies for Targeted Interventions. PLOS Computational Biology. 2014;10(8):1–20.

